## Supplemental Information for "The accuracy of absolute differential abundance analysis from relative count data"

#### Characteristics of the simulated data

The properties of simulated data sets in terms of percent differentially abundant features, change in abundance across conditions, and percent zero counts are shown in Supplemental Figure 1. Most simulations featured a minority of differentially abundant features and the distribution of the proportion of differentially abundant features was similar across feature number settings (100, 1000, and 5000 features). Fold change in total abundance between conditions was similar across settings as well. The percent zero counts in our simulations increased as the number of features increased. This is because, on average, simulations with a larger number of features had larger overall absolute total abundances and were more likely to be downsampled when ”observed” relative abundances were drawn, yielding dropouts.

Differential abundance was simulated as a random per-feature perturbation applied to a subset of randomly selected features. As such, net change in abundance across the composition was generally asymmetric. Supplemental Figure 2 illustrates the distribution of differentially abundant features experiencing an *increase* in abundance from simulated conditions A to B.

#### Visualizing outcomes from simulated data

We present alternative summaries of outcomes for all 5625 simulated data sets in this Supplement. Sensitivities and specificities organized by number of simulated features and labeled for percent of differentially abundant features are given in Supplemental Figure 3 and summaries of these outcomes are available in Supplemental Table 1. Alternatively, we present per-setting results - equivalent to those presented in main text Figure 1 - but labeled for simulated fold change in Supplemental Figure 4.

#### Variable importance in predictive models

All features utilized in predictive models are listed in Supplemental Table 2. Further, in Supplemental Tables 3 and 4, we show the top several most important features (as scored by gain in accuracy upon feature inclusion) for each of the six predictive models over sensitivity and specificity for each of ALDEx2, ANCOM-BC, DESeq2, edgeR (with TMM), and scran.

In all models over sensitivity, features summarizing the prevalence of low-count elements in the composition were highly informative. For all methods except scran, an estimates of the correlation of centered logratio features - summarizing shared change relative to the mean - were also highly informative.

Specificity was best predicted by two sets of features: 1) those describing the percent of features apparently undergoing substantial change across simulated conditions and 2) the scale of that change, as estimated by the standard deviation of observed feature-level change across conditions.

#### Results in real data

Per-method sensitivity and specificity outcomes in real data are summarized in Supplemental Tables 5 and 6 respectively. Counts of false positivies as a function of percent differentially abundant features are given in Supplemental Figure 5. See main text Figure 2 for the analogous results from simulated data.

#### Increased stringency improves agreement between absolute and relative calls

In the main text we tested for per-feature differential abundance using p-values derived from each of the methods explored after multiple test correction (via Benjamini-Hochberg) at a significance level of $\alpha \leq 0.05$. False positive counts for these results are given in main text Figure 2.

Here, we introduce a higher bar for differential abundance: a differentially abundant feature must be statistically significant with FDR $\leq 0.01$ *and* to exhibit a fold change of at least 2 (i.e. either a doubling or halving) on average across conditions to be considered truly differentially abundant. We evaluated the agreement between calls made in absolute and relative data using these criteria, in essence, asking about the degree to which data types agree of the most unambiguous differences. These results are summarized in Supplemental Figures 6 and 7. Median false positive counts for each of ANCOM-BC, DESeq2, and edgeR (with TMM) were decreased on average by 112, 101, and 102 respectively. For scran, this average FP count decrease was 54.7. For ALDEx2 were calls essentially unchanged (average change in median FP count = 0.667).

#### Specificity improvement in DESeq2 using control genes

Most of the methods we evaluated rescale the data against some per-sample measure of central tendency. An alternative to this procedure is to rescale observed abundances against an explicit reference quantity - for example, against spike-in abundances, as we have done to construct ”absolute” abundances from published data sets. DESeq2's ”control_genes” feature allows a user to specify a set of putatively stable reference features against which observed abundances can be rescaled before testing for differential abundance. We applied this control_genes feature in three ways: 1) using known stable features in simulated data, 2) using known stable features in real data, and 3) using a set of "housekeeping" genes derived from the literature [1, 2, 3, 4].

First, for each of the 5625 simulated data sets, we selected 10 features from a given data set which were at or below the 10th percentile in terms of coefficient of variation in absolute counts. These are some of the stablest features in the "true" abundances. In the corresponding relative abundance data, we identified these as control genes using the argument of the same name in DESeq2 and performed differential abundance accuracy analysis as described in the main text. The median false positive rate (FPR) across all simulated data sets was reduced from 0.124 to 0.083 and in simulated data sets with a majority of differentially abundant features, median FPR was reduced by more than 50%, from 0.385 to 0.197. Distributions of these outcomes are shown in Supplemental Figure 8 where gains in sensitivity are minimal but gains in specificity are generally substantial and positive.

In real data too, DESeq2 sensitivity and specificity were generally improved by the use of "low-variance" reference features as controls, though these improvements were often modest. These results are given in Supplemental Tables 7 and 8.

Finally, we utilized thirteen "housekeeping" genes derived from four published studies [1, 2, 3, 4]: *atp5pb*, *b2m*, *cycs*, *eer1a1*, *eef2*, *gapdh*, *hmbs*, *hprt1*, *lmna*, *ppib*, *prkg1*, *tbcb*, and *tbp*. We utilized these features as controls in six of the real data sets utilizing human or mouse expression data. These results are given in Supplemental Tables 2 and 3. Sensitivity and specificity were improved in the data sets of Song et al. and Hashimshony et al., remained largely unchanged in Muraro et al. and Hagai et al., and actually worsened in Grüen et al. and Monaco et al. This result emphasizes the need for caution in the selection of housekeeping genes as reference features, as these genes' products may be expected to exhibit variation across tissue types, organisms, and experimental settings [5].

### Supplemental Tables

| Feature number | Method | Median specificity | Percent of data sets below 95% specificity | Percent of data sets below 50% specificity |
|---|---|---|---|---|
| 100 | ALDEx2 | 0.857 | 64% | 26% |
| 100 | ANCOM-BC | 0.904 | 72% | 12% |
| 100 | DESeq2 | 0.841 | 82% | 16% |
| 100 | edgeR (TMM) | 0.903 | 71% | 15% |
| 100 | scran | 0.885 | 75% | 12% |
| 1000 | ALDEx2 | 0.925 | 58% | 9% |
| 1000 | ANCOM-BC | 0.859 | 87% | 12% |
| 1000 | DESeq2 | 0.853 | 79% | 14% |
| 1000 | edgeR (TMM) | 0.869 | 84% | 16% |
| 1000 | scran | 0.903 | 71% | 8% |
| 5000 | ALDEx2 | 0.942 | 54% | 2% |
| 5000 | ANCOM-BC | 0.822 | 87% | 14% |
| 5000 | DESeq2 | 0.879 | 73% | 9% |
| 5000 | edgeR (TMM) | 0.86 | 83% | 12% |
| 5000 | scran | 0.925 | 63% | 4% |

Table 1: Specificity in simulated data, FDR $\leq$ 0.05. Per-method results are grouped and sorted by increasing feature number.

| Feature symbol | Description |
|---|---|
| P | feature number |
| PARTIAL | totals expected to be partially informative (e.g. spike-in normalized) |
| TOTALS_C_FC | absolute fold change in mean totals (A vs. B) |
| TOTALS_C_D | absolute change in mean totals |
| TOTALS_C_MAX_D | max delta in totals |
| TOTALS_C_MED_D | median delta in totals |
| TOTALS_C_SD_D | std. dev. of in totals |
| CORR_RA_MED | median correlation of relative abundances |
| CORR_RA_SD | std. dev. of correlation of relative abundances |
| CORR_RA_SKEW | skew of correlation of relative abundances |
| CORR_LOG_MED | median correlation of log counts |

| | |
|---|---|
| CORR_LOG_SD | std. dev. of correlation of log counts |
| CORR_LOG_SKEW | skew of correlation of log counts |
| CORR_CLR_MED | median correlation of CLR features |
| CORR_CLR_SD | std. dev. of correlation of CLR features |
| CORR_CLR_SKEW | skew of correlation of CLR features |
| COMP_C_P0_A | percent features = 0 in condition A |
| COMP_C_P0_B | percent features = 0 in condition B |
| COMP_C_P1_A | percent features = 1 in condition A |
| COMP_C_P1_B | percent features = 1 in condition B |
| COMP_C_P5_A | percent features $\leq$ 5 in condition A |
| COMP_C_P5_B | percent features $\leq$ 5 in condition B |
| COMP_RA_P01_A | percent features < 0.1% relative abundance in condition A |
| COMP_RA_P01_B | percent features < 0.1% relative abundance in condition B |
| COMP_RA_P1_A | percent features < 1% relative abundance in condition A |
| COMP_RA_P1_B | percent features < 1% relative abundance in condition B |
| COMP_RA_P5_A | percent features < 5% relative abundance in condition A |
| COMP_RA_P5_B | percent features < 5% relative abundance in condition B |
| COMP_RA_MAX_A | max relative abundance in condition A |
| COMP_RA_MED_A | median relative abundance in condition A |
| COMP_RA_SD_A | std. dev. of relative abundance in condition A |
| COMP_RA_SKEW_A | skew of relative abundance in condition A |
| COMP_RA_MAX_B | max relative abundance in condition B |
| COMP_RA_MED_B | median relative abundance in condition B |
| COMP_RA_SD_B | std. dev. of relative abundance in condition B |
| COMP_RA_SKEW_B | skew of relative abundance in condition B |
| COMP_C_ENT_A | entropy in condition A |
| COMP_C_ENT_B | entropy in condition B |
| FW_RA_MAX_D | max change in relative abundance |
| FW_RA_MED_D | median change in relative abundance |
| FW_RA_SD_D | std. dev. of change in relative abundance |
| FW_RA_PPOS_D | percent features with + change in relative abundances |
| FW_RA_PNEG_D | percent features with - change in relative abundances |
| FW_RA_PFC05_D | percent features with < 0.5 FC in relative abundance |
| FW_RA_PFC1_D | percent features with < 1 FC in relative abundance |

| | |
|---|---|
| FW_RA_PFC2_D | percent features with < 2 FC in relative abundance |
| FW_LOG_MAX_D | max change in log counts |
| FW_LOG_MED_D | median change in log counts |
| FW_LOG_SD_D | std. dev. of change in log counts |
| FW_LOG_PPOS_D | percent features with + change in log counts |
| FW_LOG_PNEG_D | percent features with - change in log counts |
| FW_LOG_PFC05_D | percent features with < 0.5 FC in log counts |
| FW_LOG_PFC1_D | percent features with < 1 FC in log counts |
| FW_LOG_PFC2_D | percent features with < 2 FC in log counts |
| FW_CLR_MAX_D | max change in CLR |
| FW_CLR_MED_D | median change in CLR |
| FW_CLR_SD_D | std. dev. of change in CLR |
| FW_CLR_PPOS_D | percent features with + change in CLR |
| FW_CLR_PNEG_D | percent features with - change in CLR |
| FW_CLR_PFC05_D | percent features with < 0.5 FC in CLR |
| FW_CLR_PFC1_D | percent features with < 1 FC in CLR |
| FW_CLR_PFC2_D | percent features with < 2 FC in CLR |

Table 2: Features of observed abundances used to predict sensitivity and specificity outcomes.

| Method | Feature | Importance |
|---|---|---|
| ALDEx2 | percent features = 0 in condition B | 1 |
| ALDEx2 | skew of correlation of CLR features | 0.78 |
| ALDEx2 | median correlation of CLR features | 0.51 |
| ANCOM-BC | percent features = 1 in condition A | 1 |
| ANCOM-BC | percent features $\leq$ 5 in condition A | 0.82 |
| ANCOM-BC | median correlation of CLR features | 0.79 |
| DESeq2 | percent features $\leq$ 5 in condition B | 1 |
| DESeq2 | median correlation of CLR features | 0.94 |
| DESeq2 | percent features = 1 in condition B | 0.74 |
| edgeR (TMM) | median correlation of CLR features | 1 |
| edgeR (TMM) | skew of correlation of CLR features | 0.74 |
| edgeR (TMM) | percent features = 0 in condition B | 0.71 |
| scran | percent features $\leq$ 5 in condition B | 1 |
| scran | percent features = 1 in condition A | 0.95 |
| scran | percent features $\leq$ 5 in condition A | 0.94 |

Table 3: Predictive features and their relative importance (as gain) in the prediction of sensitivity.

| Method | Feature | Importance |
|---|---|---|
| ALDEx2 | percent features with $<$ 0.5 FC in CLR | 1 |
| ALDEx2 | std. dev. of change in log counts | 0.45 |
| ALDEx2 | percent features = 0 in condition B | 0.39 |
| ANCOM-BC | percent features with $<$ 0.5 FC in CLR | 1 |
| ANCOM-BC | percent features with $<$ 0.5 FC in log counts | 0.39 |
| ANCOM-BC | std. dev. of change in log counts | 0.37 |
| DESeq2 | percent features with $<$ 0.5 FC in CLR | 1 |
| DESeq2 | std. dev. of change in log counts | 0.44 |
| DESeq2 | percent features with $<$ 2 FC in CLR | 0.42 |
| edgeR (TMM) | percent features with $<$ 0.5 FC in CLR | 1 |
| edgeR (TMM) | percent features with $<$ 2 FC in CLR | 0.44 |
| edgeR (TMM) | std. dev. of change in log counts | 0.36 |
| scran | percent features with $<$ 0.5 FC in CLR | 1 |
| scran | std. dev. of change in log counts | 0.51 |
| scran | percent features with $<$ 2 FC in CLR | 0.34 |

Table 4: Predictive features and their relative importance (as gain) in the prediction of specificity.

| Data set | ALDEx2 | ANCOM-BC | DESeq2 | edgeR (TMM) | scran |
|---|---|---|---|---|---|
| Hagai et al. | 0.087 | 0.574 | 0.465 | 0.521 | 0.574 |
| Hashimshony et al. | 0 | 0.093 | 0.104 | 0.134 | 0.072 |
| Song et al. | 0.398 | 0.472 | 0.472 | 0.5 | 0.535 |
| Monaco et al. | 0.056 | 0.449 | 0.373 | 0.391 | 0 |
| Vieira-Silva et al. | 0 | 0.444 | 0.528 | 0.556 | 0.139 |
| Barlow et al. | 0 | 0.082 | 0 | 0.061 | 0.122 |
| Gruen et al. | 0.229 | 0.436 | 0.748 | 0.604 | 0.536 |
| Muraro et al. | 0.128 | 0.334 | 0.316 | 0.415 | 0.371 |
| Kimmerling et al. | 0.004 | 0.012 | 0.002 | 0.005 | 0.084 |
| Yu et al. | 0.826 | 0.905 | 0.896 | 0.889 | 0.898 |
| Owens et al. | 0.741 | 0.835 | 0.836 | 0.833 | 0.823 |
| Klein et al. | 0.397 | 0.66 | 0.645 | 0.607 | 0.654 |

Table 5: Observed sensitivities on real data sets.

| Data set | ALDEx2 | ANCOM-BC | DESeq2 | edgeR (TMM) | scran |
|---|---|---|---|---|---|
| Hagai et al. | 0.983 | 0.825 | 0.968 | 0.96 | 0.862 |
| Hashimshony et al. | 1 | 0.987 | 0.972 | 0.967 | 0.99 |
| Song et al. | 0.771 | 0.624 | 0.653 | 0.719 | 0.865 |
| Monaco et al. | 0.991 | 0.634 | 0.83 | 0.77 | 1 |
| Vieira-Silva et al. | 0.975 | 0.75 | 0.95 | 0.875 | 0.925 |
| Barlow et al. | 1 | 0.927 | 0.909 | 0.855 | 1 |
| Gruen et al. | 0.913 | 0.671 | 0.921 | 0.878 | 0.794 |
| Muraro et al. | 0.97 | 0.778 | 0.853 | 0.904 | 0.834 |
| Kimmerling et al. | 1 | 0.994 | 0.998 | 0.993 | 0.993 |
| Yu et al. | 0.773 | 0.56 | 0.855 | 0.754 | 0.536 |
| Owens et al. | 0.147 | 0.064 | 0.082 | 0.04 | 0.147 |
| Klein et al. | 0.036 | 0.018 | 0.089 | 0.018 | 0.018 |

Table 6: Observed specificities on real data sets.

| Data set | No controls | Low-variance features | Housekeeping genes |
|---|---|---|---|
| Song | 0.472 | 0.947 | 0.866 |
| Monaco | 0.373 | 0.454 | 0.432 |
| Muraro | 0.316 | 0.391 | 0.315 |
| Hagai | 0.465 | 0.5 | 0.476 |
| Hashimshony | 0.104 | 0.201 | 0.253 |
| Gruen | 0.748 | 0.785 | 0.447 |

Table 7: DESeq2 sensitivity when using a variety of features as controls (via the control_genes argument). Sensitivity is improved when using a random set of low-variance features as references against which to rescale observed abundances. A selection of "housekeeping" genes used for the same purpose gives less improvement over baseline.

| Data set | No controls | Low-variance features | Housekeeping genes |
|---|---|---|---|
| Song | 0.653 | 0.902 | 0.886 |
| Monaco | 0.83 | 0.896 | 0.575 |
| Muraro | 0.853 | 0.906 | 0.872 |
| Hagai | 0.968 | 0.975 | 0.97 |
| Hashimshony | 0.972 | 0.986 | 0.989 |
| Gruen | 0.921 | 0.924 | 0.748 |

Table 8: DESeq2 specificity when using a variety of features as controls (via the control_genes argument). Specificity is improved when using a random set of low-variance features as references against which to rescale observed abundances. A selection of "housekeeping" genes used for the same purpose gives less improvement or worsens specificity in the data sets of Monaco et al. and Grüen et al.

#### Supplemental Figures

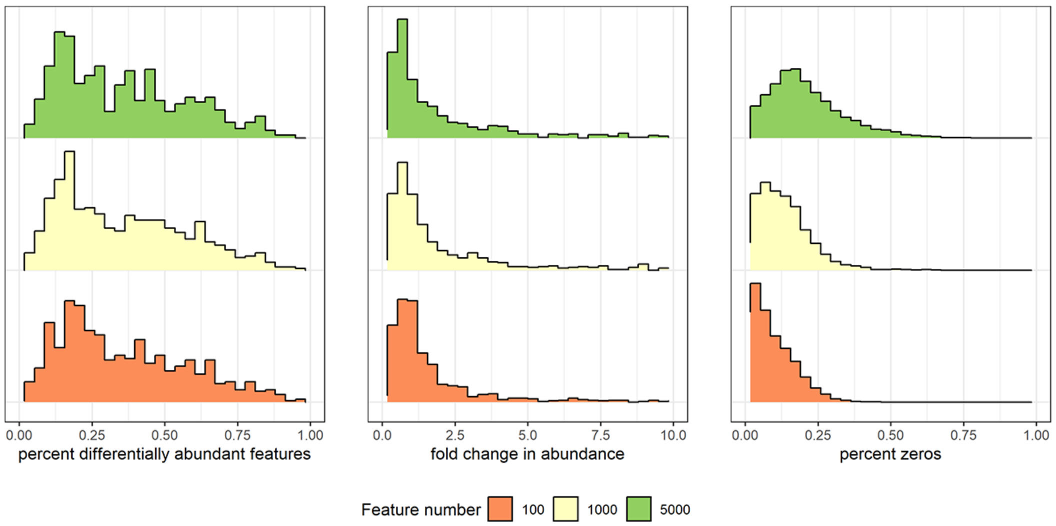

Figure 1: Distributions associated with three characteristics of the 5625 simulated data sets: **a)** percent differentially abundant features, **b)** fold change in total abundance across conditions, and **c)** percent zeros.

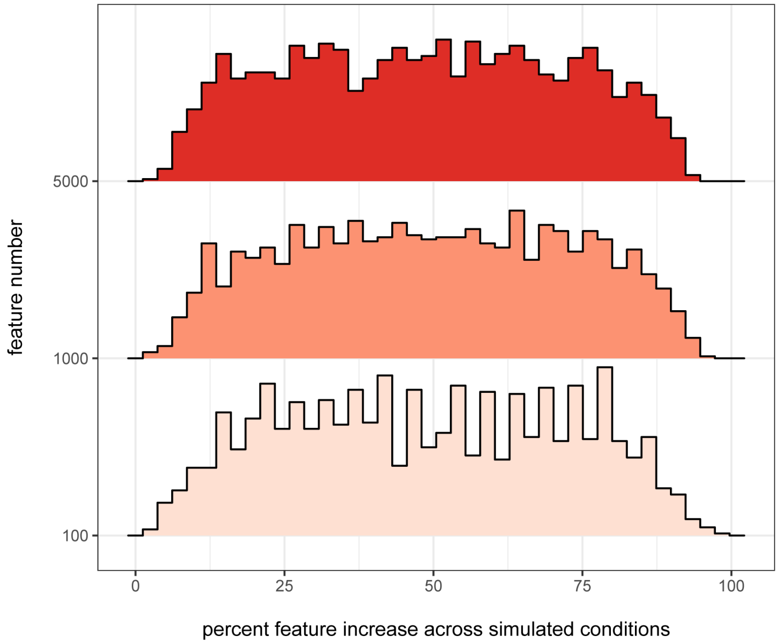

Figure 2: The number, scale, and direction of differential abundance varied randomly in simulation, giving rise to both increases and decreases in total abundance across conditions.

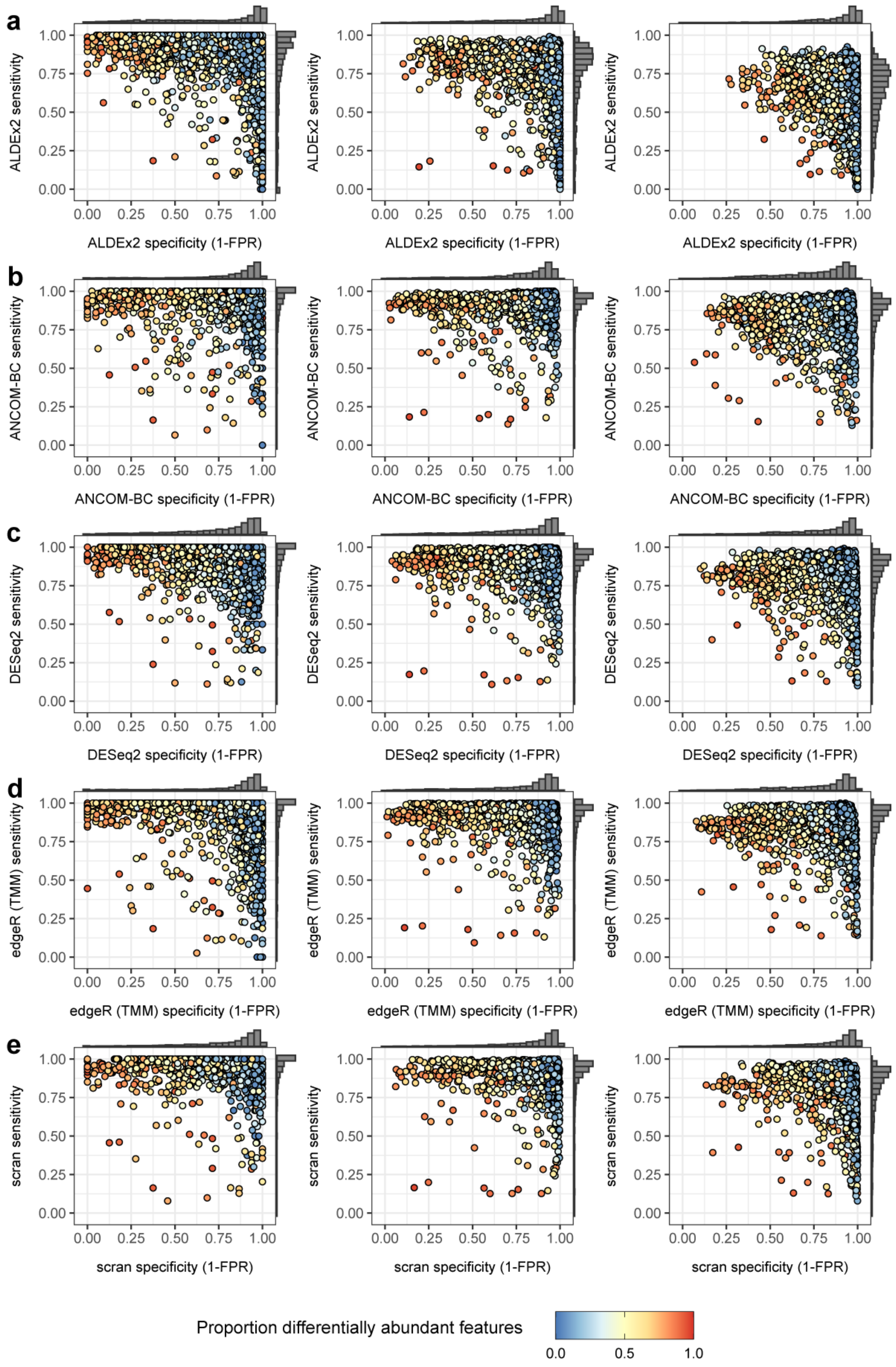

Figure 3: Sensitivity and specificity for five differential abundance calling methods in terms of increasing feature number from **a)** 100 to **b)** 1000 to **c)** 5000 features.

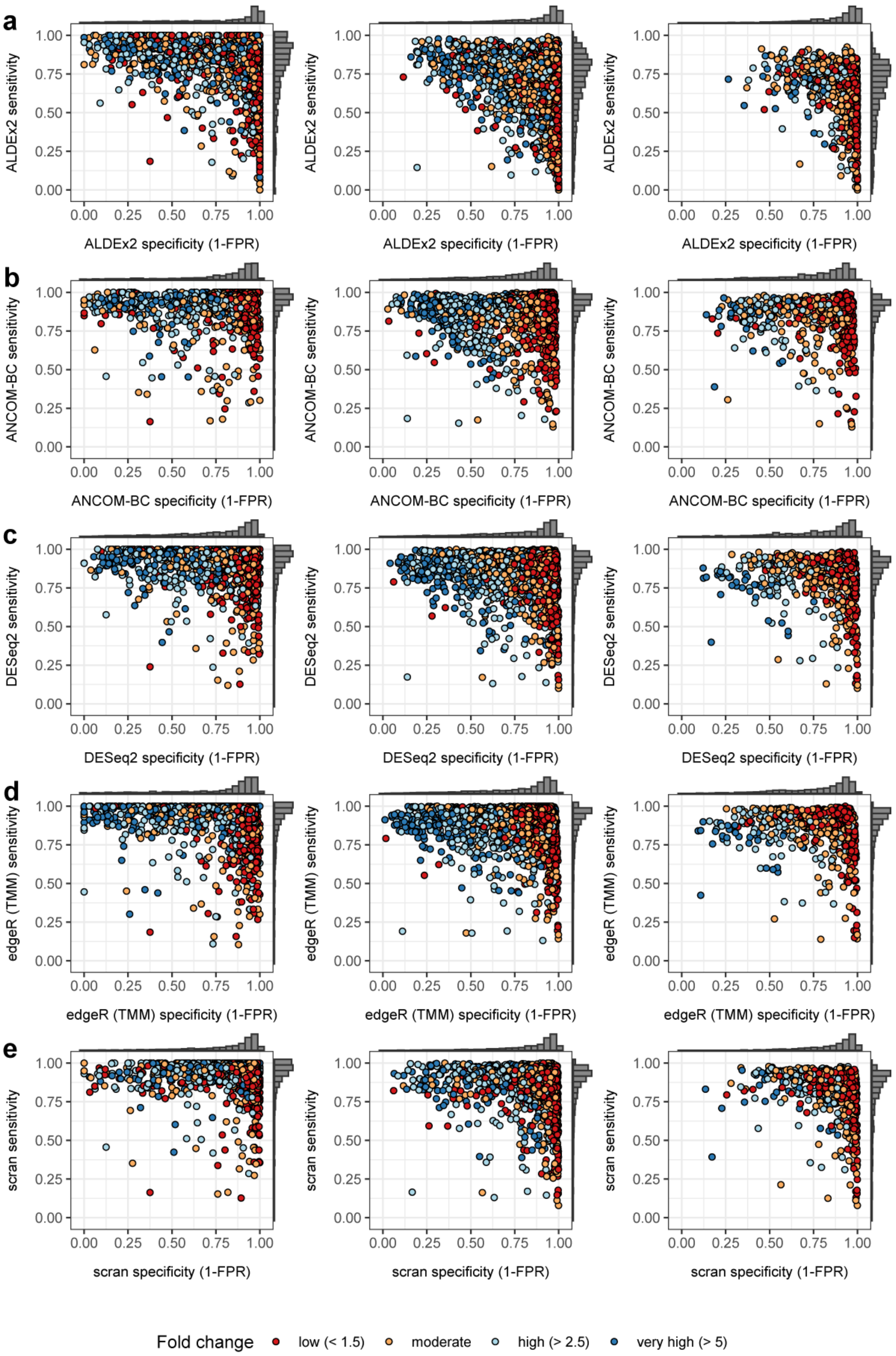

Figure 4: Sensitivity and specificity for five differential abundance calling methods in three experimental settings: **a)** Microbial, **b)** Bulk Transcriptomic, and **c)** Cell Transcriptomic settings. Data sets are labeled by fold change across conditions.

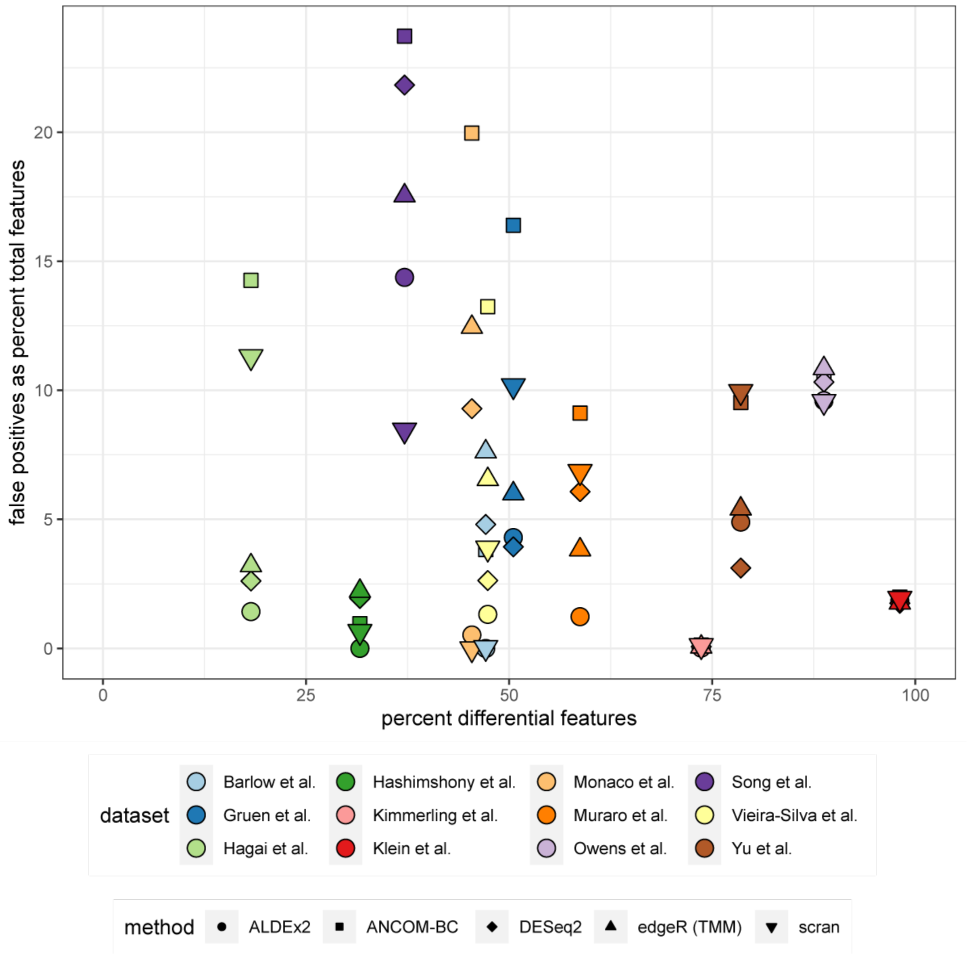

Figure 5: Counts of false positives as a function of the percent of simulated differentially abundant features for five methods applied to 12 real data sets.

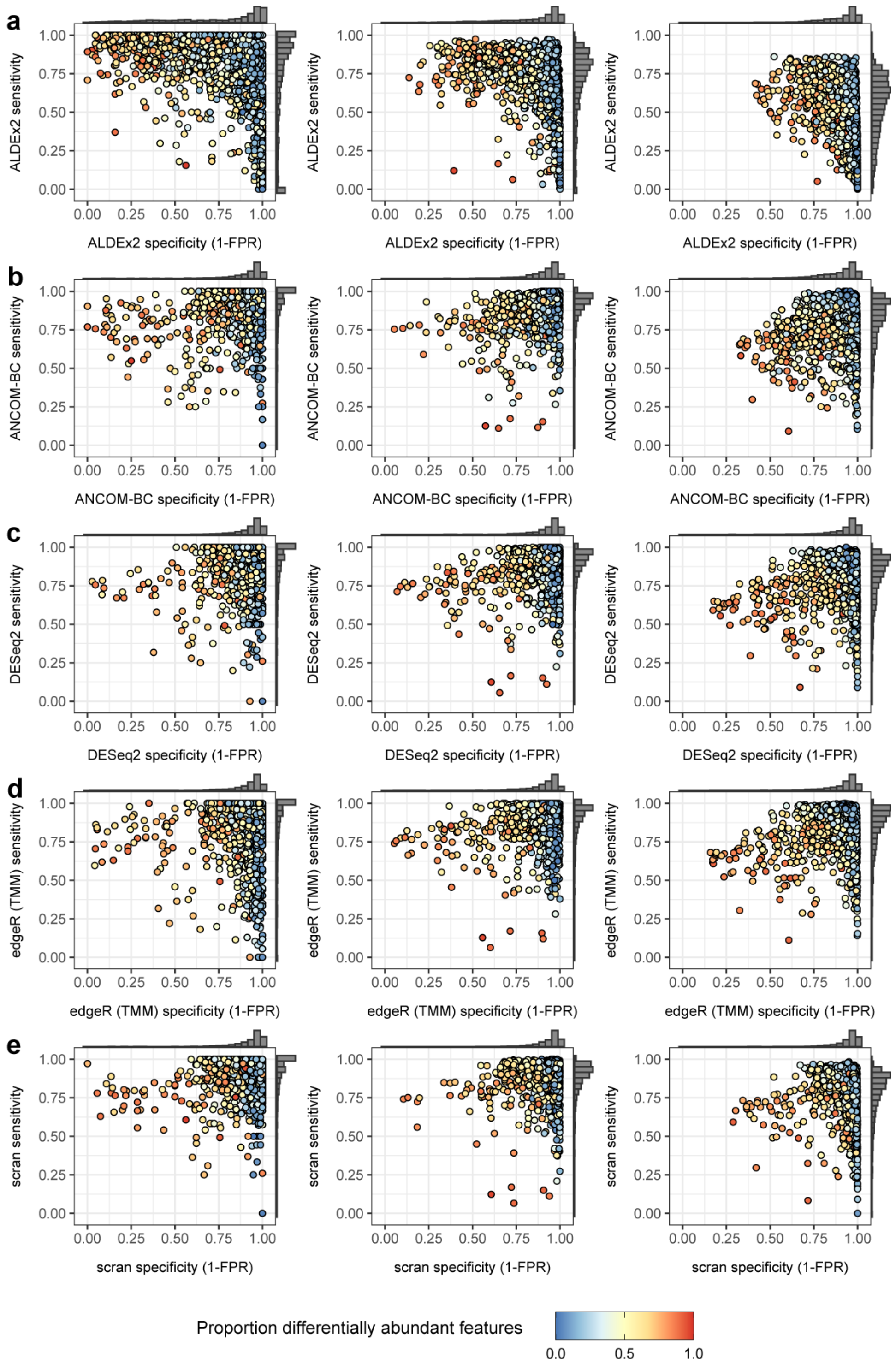

Figure 6: Sensitivity and specificity for five differential abundance calling methods in three experimental settings: **a)** Microbial, **b)** Bulk Transcriptomic, and **c)** Cell Transcriptomic settings. Data sets are labeled by proportion of differentially abundant features. Here, differential abundance calling is subject to greater stringency: FDR $\leq 0.01$ and a fold change across conditions of at least 2. Median specificity is improved from 0.90 (FDR $\leq 0.05$) to 0.95.

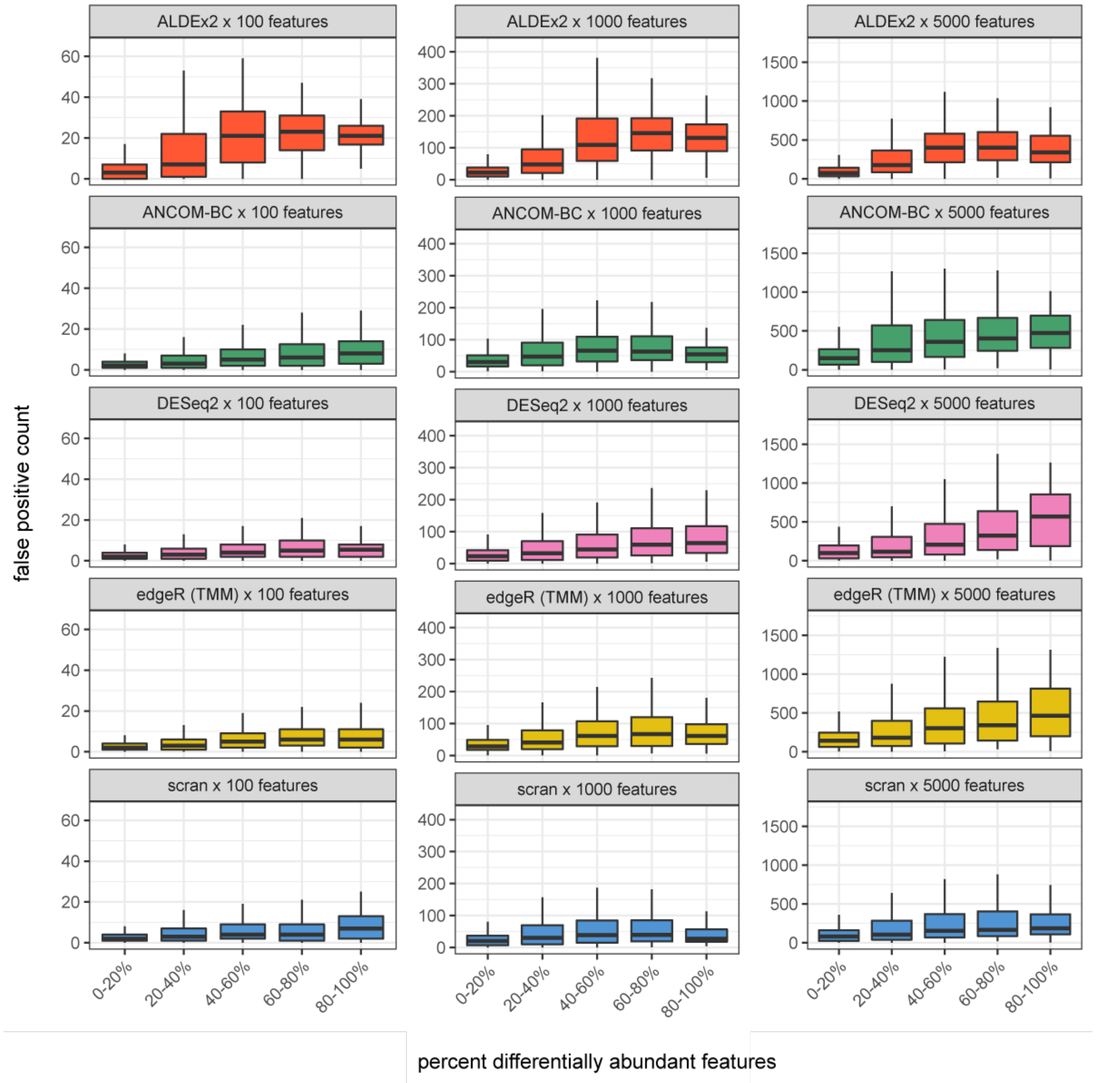

Figure 7: Counts of false positives as a function of the percent of simulated differentially abundant features for five methods with stringent differential abundance calling (FDR $\leq$ 0.01 and a fold change across conditions of at least 2). Columns segregate simulations with increasing numbers of features. Counts of false positives are reduced relative to FDR $\leq$ 0.05.

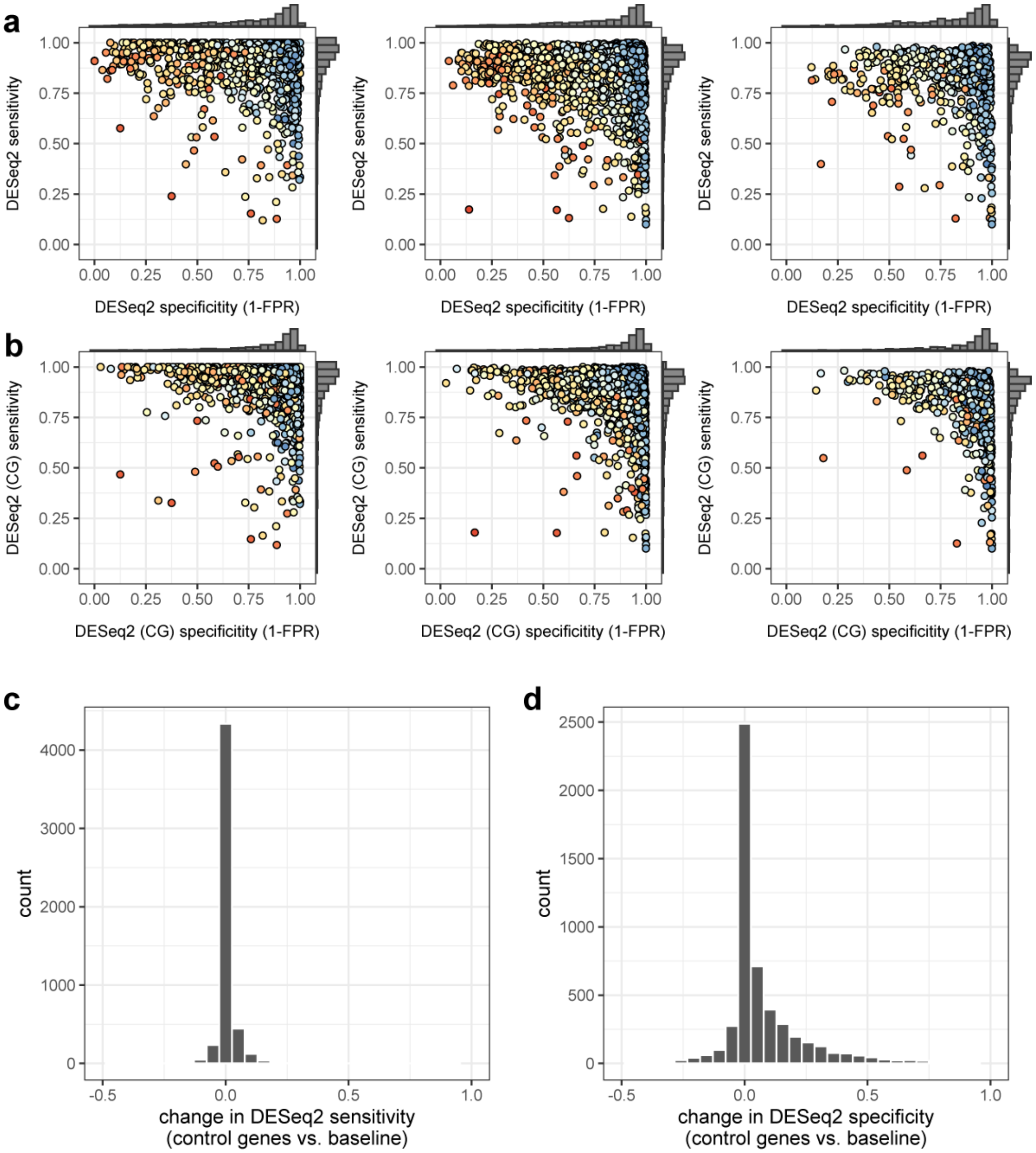

Figure 8: Sensitivity and specificity for all simulated data sets using DESeq2 **a)** without and **b)** with "control genes." Per-columns settings from left to right are Microbial, Transcriptomic (center column), and Bulk Transcriptomic. **c)** Distribution of the change in sensitivity and **d)** specificity following the introduction of control genes. Sensitivity is largely unchanged. Specificity is generally improved.
